# Discovery of interaction-related sRNAs and their targets in the *Brachypodium distachyon* and *Magnaporthe oryzae* pathosystem

**DOI:** 10.1101/631945

**Authors:** Silvia Zanini, Ena Šečić, Tobias Busche, Jörn Kalinowski, Karl-Heinz Kogel

## Abstract

Microbial pathogens secrete small RNA (sRNA) effectors into plant hosts to aid infection by silencing transcripts of immunity and signaling-related genes through RNA interference (RNAi). Similarly, sRNAs from plant hosts have been shown to contribute to plant defense against microbial pathogens by targeting transcripts involved in virulence. This phenomenon is called bidirectional RNA communication or cross kingdom RNAi (ckRNAi). How far this RNAi-mediated mechanism is evolutionarily conserved is the subject of controversial discussions. We examined the bidirectional accumulation of sRNAs in the interaction of the hemibiotrophic rice blast fungus *Magnaporthe oryzae* (*Mo*) with the grass model plant *Brachypodium distachyon* (*Bd*). By comparative deep sequencing of sRNAs and mRNAs from axenic fungal cultures and infected leaves and roots, we found a wide range of fungal sRNAs that accumulated exclusively in infected tissues. Amongst those, 20-21 nt candidate sRNA effectors were predicted *in silico* by selecting those *Mo* reads that had complementary mRNA targets in *Bd*. Many of those mRNAs predicted to be targeted by *Mo* sRNAs were differentially expressed, particularly in the necrotrophic infection phase, including gene transcripts involved in plant defense responses and signaling. Vice versa, by applying the same strategy to identify *Bd* sRNA effectors, we found that *Bd* produced sRNAs targeting a variety of fungal transcripts, encoding fungal cell wall components, virulence genes and transcription factors. Consistent with function as effectors of these *Bd* sRNAs, their predicted fungal targets were significantly down-regulated in the infected tissues compared to axenic cultures, and deletion mutants for some of these target genes showed heavily impaired virulence phenotypes. Overall, this study provides the first experimentally-based evidence for bidirectional ckRNAi in a grass-fungal pathosystem, paving the way for further validation of identified sRNA-target duplexes and contributing to the emerging research on naturally occurring cross-kingdom communication and its implications for agriculture on staple crops.

**Author Summary:** In the present work, we provide first experimental evidence for bidirectional RNA communication in a grass-fungal pathosystem. We deployed the monocotyledonous plant *Brachypodium distachyon*, which is a genetic model for the staple crops wheat and rice, to investigate the interaction-related sRNAs for their role in RNA communication. By applying a previously published bioinformatics pipeline for the detection of sRNA effectors we identified potential plant targets for fungal sRNAs and vice versa, fungal targets for plant sRNAs. Inspection of the respective targets confirmed their downregulation in infected relative to uninfected tissues and fungal axenic cultures, respectively. By focusing on potential fungal targets, we identified several genes encoding fungal cell wall components, virulence proteins and transcription factors. The deletion of those fungal targets has already been shown to produce disordered virulence phenotypes. Our findings establish the basis for further validation of identified sRNA-mRNA target duplexes and contribute to the emerging research on naturally occurring cross-kingdom communication and its implications for agriculture.

## Introduction

Small (s)RNAs such as small interfering (si)RNAs, micro (mi)RNAs, and transfer (t)RNAs are systemic signals that modulate distal gene regulation and epigenetic events in response to biotic and abiotic environmental cues in plants (Molnar et al. 2010 Borges & Martienssen 2015; Kehr & Kragler 2018). Particularly, sRNA-mediated gene silencing is one of the main defense mechanisms against viral attack and damaging effects of transposons. The action of sRNAs rests upon their role in RNA interference (RNAi), a conserved mechanism of gene regulation in eukaryotes at the translational (PTGS or post-transcriptional gene silencing) and transcriptional (TGS or transcriptional gene silencing) level (Fire et al. 1998; Vaucheret & Fagard 2001; Castel & Martienssen 2013). In plants, the trigger for RNAi is either endogenous or exogenous (e.g. viral) double-stranded (ds)RNA that is cut into short 20 to 24 nucleotide (nt) sRNA by DICER-like (DCL) enzymes (Hamilton & Baulcombe 1999; Baulcombe 2004). The duplexes are incorporated into an RNA-induced silencing complex (RISC), containing an endonucleolytic ARGONAUTE (AGO) protein to target partially complementary RNAs for mRNA degradation/inhibition or epigenetic modification by RNA-dependent DNA methylation (RdDM), histone modification and chromatin remodeling, while plant RNA-dependent RNA polymerases (RdRPs) are involved in the production of secondary sRNAs (Castel & Martienssen, 2013; Vaucheret et al. 2004).

Consistent with the movement of RNAs during animal-parasitic interactions (Buck et al. 2014; LaMonte et al. 2012; Garcia-Silva et al. 2014), recent reports suggest that sRNAs also move from plants into fungal pathogens and, vice versa, from pathogens to plants to positively or negatively regulate genes involved in pathogenesis (Weiberg et al. 2013; Zhang et al. 2016; Wang et al. 2017a; Wang et al. 2017b). First hints for this “bidirectional” or “cross kingdom” RNAi (ckRNAi) and the action of sRNA effectors in plants originally came from studies that showed efficient delivery of artificially designed sRNA from plants into interacting microbes. Such plant-mediated RNAi, termed host-induced gene silencing (HIGS, Nowara et al. 2010), includes formation of dsRNA from hairpin or inverted promoter constructs, dsRNA processing into sRNAs and transfer of these into the interacting microbe. As of today, HIGS has emerged as a promising strategy for crop protection against viruses, fungi, oomycetes, nematodes, and insects (Head et al. 2017; Koch et al. 2013; Govindarajulu et al. 2015; for review see Cai et al. 2018a). The broad applicability of the biotechnological HIGS technique implied the possibility of an evolutionarily-conserved mechanism of sRNA cross-kingdom trafficking. Consistent with this view, the plant-pathogenic fungus *Verticillium dahliae* (*Vd*) recovered from infected cotton plants, contained plant miRNAs, implying that host-derived sRNAs were transmitted into the pathogen during infection (Zhang et al. 2016). Two of those cotton miRNAs, miR166 and miR159, target the fungal genes *Ca^2+^-DEPENDENT CYSTEINE PROTEASE CALPAIN* (*VdClp-1*) and *ISOTRICHODERMIN C-15 HYDROXYLASE* (*VdHiC-15*), respectively, which are known to contribute to fungal virulence.

Similarly, *Arabidopsis* cells secrete vesicles to deliver sRNAs into grey mold fungal pathogen *Botrytis cinerea* (Cai et al. 2018b). These sRNA-containing vesicles accumulate at the infection sites and are taken up by the fungal cells to induce silencing of fungal genes critical for its pathogenicity. Consistent with the bidirectional move of sRNAs in plant-microbe interactions, *B. cinerea* also produces sRNA effectors, predicted to originate from long-terminal repeat (LTR) retrotransposons in the fungal genome, that down-regulate *Arabidopsis* and tomato genes involved in immunity (Weiberg et al. 2013). Some of those sRNA effectors were shown to target a large set of host immunity genes to enhance *B. cinerea* (*Bc*) pathogenicity, for example *Bc*-siR37, able to suppress the plant host immunity by targeting various *Arabidopsis* genes, including WRKY transcription factors, receptor-like kinases, and cell wall-modifying enzymes (Wang et al. 2017b).

The mechanism of sRNAs transfer in plant host - microbe interactions is proposed to be via plant extracellular vesicles (EVs), derived from multivesicular bodies (MVBs; An et al. 2006a, 2006b) form lipid compartments capable of trafficking proteins, lipids, and metabolites between cells, and were shown to be enriched in stress response proteins and signaling lipids and displayed antifungal activity (Rutter and Innes 2017). Consistent with the work on animal EVs (Buck et al. 2014), plant EVs also contain sRNAs such as miRNAs, tasiRNAs and heterochromatic sRNAs derived from intergenic regions (Cai et al. 2018b; Baldrich et al. 2019).

Because only a few studies have been published since the landmark paper of Weiberg et al. (2013), the extent of occurrence of sRNA effectors in host-microbe interactions is unclear and their involvement is even challenged for certain pathosystems (Kettles et al. 2018). Here we investigate the potential cross-kingdom role of sRNAs in the interaction of *Magnaporthe oryzae* (*Mo*) with *Brachypodium distachyon* (*Bd*). *Mo* is a hemibiotrophic fungal pathogen causing rice blast, the most devastating disease of cultivated rice, and is of global economic importance (Dean et al. 2012; Donofrio et al. 2014). The fungus also infects other cereals, including barley, rye, and wheat, making it a major threat to global food security (Sesma & Osbourn 2004; Wilson and Talbot 2009). *Mo* infections also have been established in the grass *B. distachyon* (Routledge et al. 2004; Parker et al. 2008). *Bd* is preferable to more complex crops, such as hexaploid wheat (*Triticum aestivum*), with a fully sequenced genome due to its smaller genome size (272 Mb) and complexity, a short life cycle and a vast T-DNA insertion library available (Fitzgerald et al. 2015; Vogel et al. 2006).

Expression of endogenous sRNAs in *Bd* following abiotic stress has been shown, pointing to operable RNAi-based regulatory mechanisms in this plant species (Wang et al. 2015). Major components of *Bd*’s RNAi machinery have been identified *in silico*, resulting in 16 BdAGO-like and six BdDCL candidates (Mirzaei et al. 2014; Secic et al. 2019). The genome of *Mo* encodes for two *DCL* genes, three *AGO* genes, and three *RdRP* genes (Kadotani et al. 2003; Murphy et al. 2008; Raman et al. 2017). According to a recent publication, *MoDCL2*, but not *MoDCL1*, is necessary for siRNA production from dsRNA (Raman et al. 2017). The analysis of sRNA in *Mo* has identified methylguanosine-capped and polyadenylated sRNA (Gowda et al. 2010) as well as sRNA matching repeats, intergenic regions (IGR), transfer RNA (tRNA), ribosomal RNA (rRNA), small nuclear (snRNA), and protein-coding genes (Nunes et al. 2011; Raman et al. 2013). Mutations in *MoDCL2*, *MoRdRP2*, and *MoAGO3* reduced sRNA levels (Raman et al. 2017), suggesting that MoDCL2, MoRdRP2 and MoAGO3 are required for the biogenesis and function of sRNA-matching genome-wide sites such as coding, intergenic regions and repeats. Of note, loss of MoAGO3 function reduced both sRNAs and fungal virulence on barley leaves. Moreover, transcriptome analysis of multiple *Mo* mutants revealed that sRNAs play an important role in transcriptional regulation of repeats and intergenic regions (Raman et al. 2017). Taken together, these findings support the notion that *Mo* sRNAs regulate fungal developmental processes, including growth and virulence. Here we further explore the role of *Mo* and *Bd* sRNAs in the *Mo*-*Bd* interaction based on data generated by parallel sRNA and mRNA deep sequencing of infected leaf and root material. Following a previously published bioinformatics pipeline (Zanini et al. 2018) for characterization of sRNA effectors and their targets, we found strong evidence for ckRNAi in a grass pathosystems.

## Results

### Selection of interaction-related sRNAs in the *Mo-Bd* pathosystem

To establish a ckRNAi function of sRNAs in the interaction between *Magnaporthe oryzae* (*Mo* 70-15) and *Brachypodium distachyon* (*Bd21-3*), we first isolated sRNA and mRNA fractions of total RNA from the same biological material (roots, leaves and axenic mycelium) and after cDNA library preparation performed high throughput next generation sequencing (NGS). TruSeq^®^ Small RNA libraries and TruSeq® Stranded mRNA libraries were produced from *i. Mo* axenic culture, *ii. Mo*-infected and mock-treated *Bd* roots (at 4 DPI), and *iii. Mo*-infected and mock-treated *Bd* leaves (at 2 DPI and 4 DPI) (Fig. 1). These time points were chosen to cover both the biotrophic and necrotrophic phase of leaf infections of the hemibiotrophic *Mo* (Wilson and Talbot 2009). Before sequencing, multiplexed sRNA libraries were size selected for 15 to 35 nt (140-160 nt including TruSeq adapters). Single end sequencing on Illumina HiSeq1500 platform generated between 22 million (mil) and 38 mil reads each (S1 Tab). Reads were further processed and filtered based on our previously published pipeline (Zanini et al. 2018). Quality check of raw reads was performed with FastQC, adapters were removed with cutadapt and the organism of origin of the trimmed reads was predicted by mapping via Bowtie alignments to both *Bd* and *Mo* genomes (Zerbino et al. 2018, Bd21-3 v1.1 DOE-JGI, http://phytozome.jgi.doe.gov/). Ambiguous reads that could not be assigned to the organism of origin with high confidence were excluded to avoid miscalling. As expected, most reads in *Mo*-infected plant samples were assigned to *Bd* (with 100% match) and not to the fungus (with at least two nucleotide mismatches) (S1 Tab). Size distribution of genome matched unique sRNA reads followed a similar trend throughout samples, with the *Mo* reads showing a peak between 19-21 nt and *Bd* reads at 24 nt (Fig 2A-2B, S1A-S1B Fig.). In order to further investigate the sRNAs potentially playing a role in the *Mo-Bd* interaction, fungal unique sRNA reads were compared among samples from *Mo* axenic culture and *Mo*-infected leaves and roots and classified as shared or exclusive between samples (Fig 3A). Some 5,708 *Mo* sRNAs were identified in *Bd*-infected roots tissue of which 3,263 (57.15%) were found exclusively in the infected sample and not in the axenic culture. Moreover, 7,215 *Mo* sRNAs (exclusively found in infected samples: 4,399 [60.97%]) were identified in *Bd*-infected leaf tissue during the biotrophic phase and 63,017 (exclusively found in infected samples: 46,212 [73.33%]) in *Bd*-infected leaf tissue during the necrotrophic phase.

**Figure 1.**
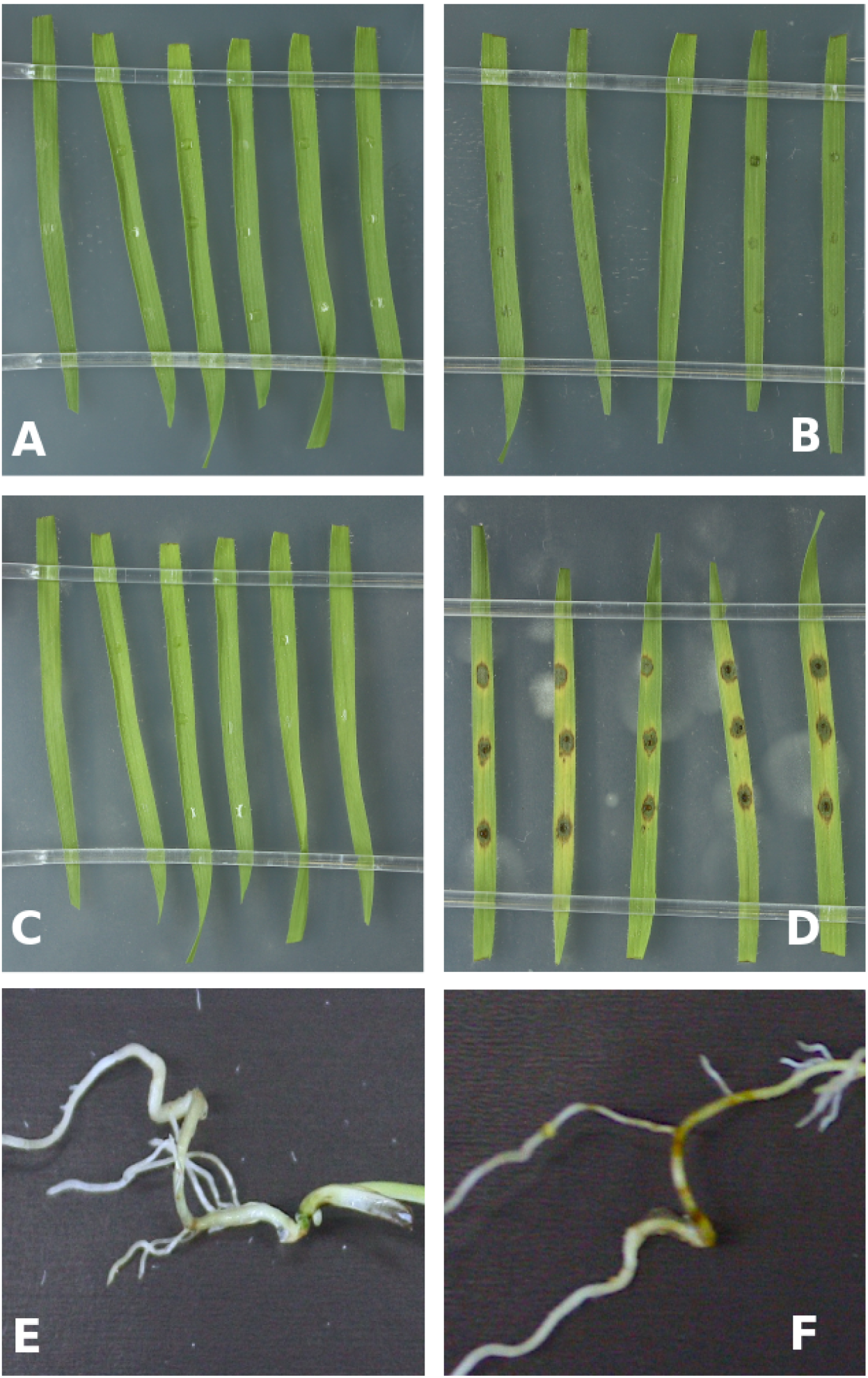
The interaction of *Brachypodium distachyon* and *Magnaporthe oryzae*. (**A,D**) Detached 21-day-old *Bd21-3* leaves were drop-inoculated with 10 μl of *Mo* 70-15 conidia solution (50,000 conidia/ml in 0.002% Tween water) and kept for 2 days (B) and 4 days (D), respectively, at high humidity. Respective controls were mock-inoculated (A,C). **(E,F)** Roots of seven-day-old seedlings were inoculated with 1 ml of conidia solution (250,000 conidia/ml in 0.002% Tween water)and kept for four days under high humidity at 16 h light/8 h dark cycle at 22°C/18°C (F). Mock-treated roots served as control (E).

**Figure 2.**
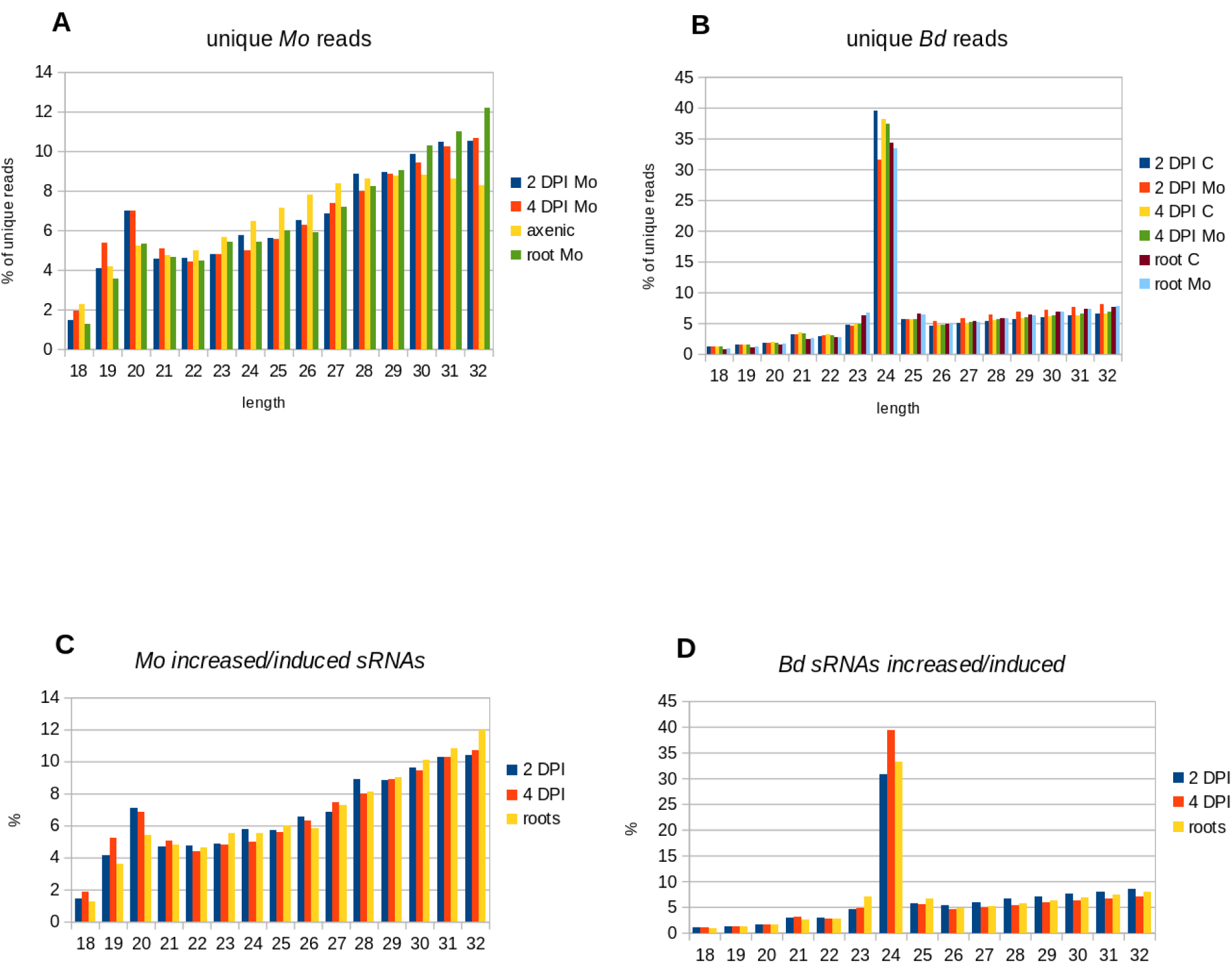
Size distribution of unique sRNA reads in the interaction of *Brachypodium distachyon* and *Magnaporthe oryzae.* (**A,B**) Relative size distribution (in percentage) of unique filtered sRNA reads assigned to *Mo* (A) or *Bd* (B) in the interaction of *M. oryzae* (*Mo* 70-15) and *B. distachyon* (*Bd21-3*). Reads were assigned to either *Mo* or *Bd* only if aligning 100% to the organism of origin genome and had at least two mismatches to the interacting organism genome. **(C,D)** Relative size distribution of unique filtered sRNA reads assigned to *Mo* (C) or *Bd* (D) and induced or increased in infected samples compared to controls (axenic fungal cultures and non-inoculated plants, respectively). Samples for sRNA sequencing by Illumina HiSeq1500 were taken from different setups: leaves (leaf 2 DPI, 4 DPI) and roots (roots 4 DPI).

**Figure 3.**
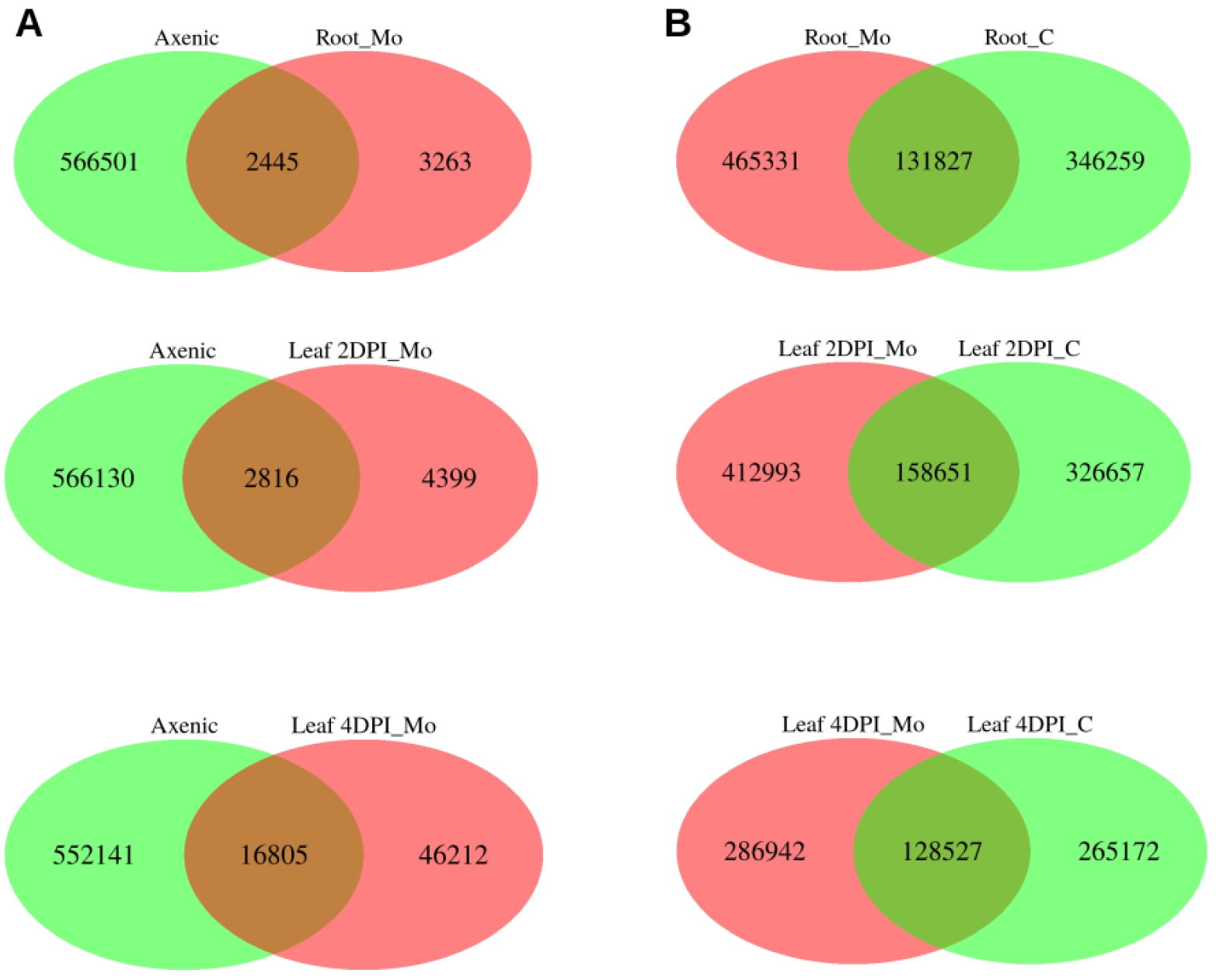
Venn diagrams of unique filtered *Mo* and *Bd* reads. (**A**) Venn diagram of *Mo* sRNA reads (18-32 nt) in axenic culture (green) and *Mo*-infected (red) *Bd* leaves (2 DPI, 4 DPI) and roots (4 DPI). (**B**) Venn diagram of *Bd* sRNA reads (18-32 nt) in mock-inoculated, non-infected (green) and *Mo*-infected (red) *Bd* leaves.

Equally, unique *Bd* sRNA reads were compared in root and leaf samples from *Mo*-infected and mock-treated *Bd21-3* (Fig 3B). We found a huge number of *Bd* sRNAs in *Mo*-infected plant tissues: 597,158 *Bd* sRNAs in *Mo*-infected roots of which 346,259 (77.92%) were solely found in infected, but not in non-infected roots; 571,644 in leaves during biotrophic interaction (2 DPI) of which 326,657 (72.24%) were solely found in infected leaves; and 415,469 during the necrotrophic interaction (4 DPI) of which 265,172 (69.06%) were solely found in infected leaves. This data suggests that most unique sRNAs from both interacting organisms are expressed exclusively during the interaction and thus are potentially of high relevance for the outcome of the disease. We selected unique sRNAs that *i.* were either found exclusively in infected plant tissues or *ii.* showed higher numbers in the infected tissue as compared to mock-infected tissue and axenic culture. Interestingly, the size distribution of these induced sRNA reads did not highlight a change in length preference compared to the total unique reads (Fig 2C-2D).

Given that ckRNAi in plant host-pathogen interactions would require an operable RNAi pathway (Weiberg et al. 2013), we tested available *Mo* mutants that are compromised for DCL and AGO activities. As shown in Fig. 4, all mutants showed reduced virulence and infection phenotypes were clearly distinguishable from the *Mo* 70-15 wild type. Of note, *Mo* Δ*dcl1* produced smaller lesions than Δ*dcl2*, suggesting that MoDCL1 plays a critical role in the *Bd-Mo* interactions. Consistent with this notion, the double mutant Δ*dcl1* Δ*dcl2* produced similar lesions to Δ*dcl1*.

**Figure 4.**
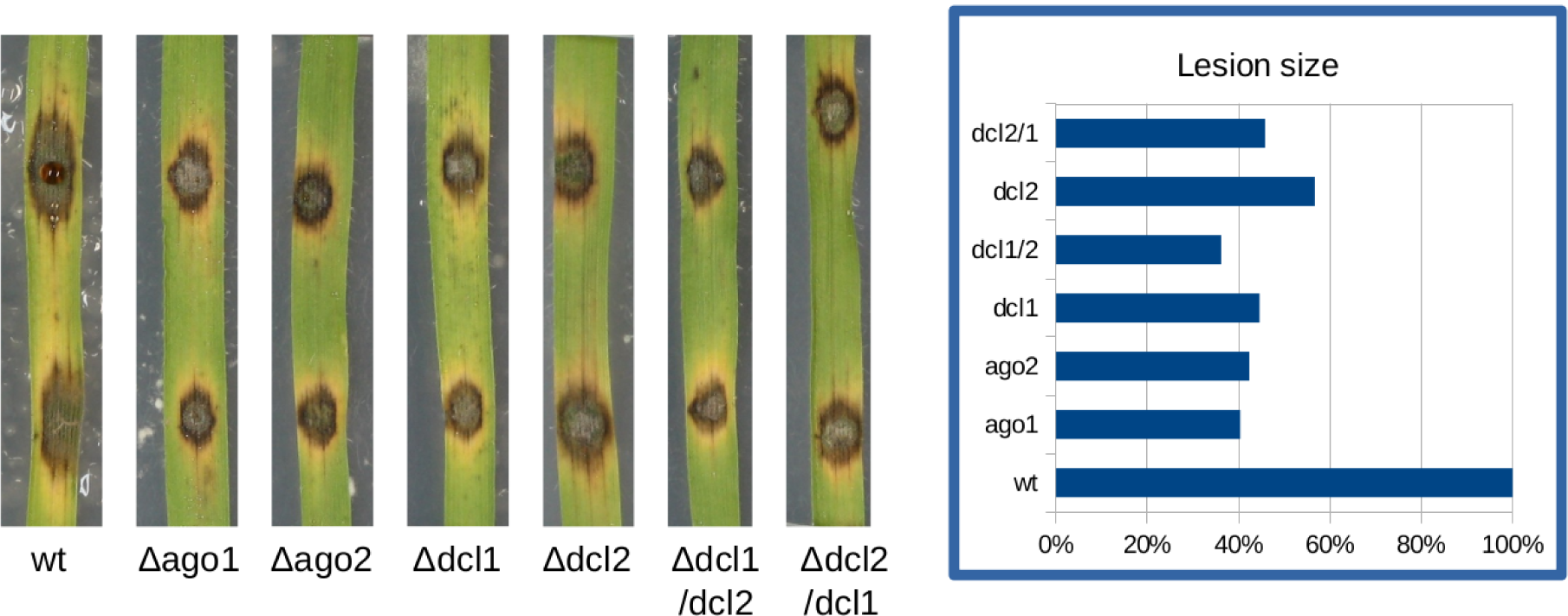
Infection phenotypes of *Magnaporthe oryzae* RNAi mutants on *Brachypodium distachyon* leaves. Detached 21-day-old *Bd21-3* leaves were drop-inoculated with 10 μl of *Mo* 70-15 conidia (50,000 conidia/ml in 0.002% Tween water) and kept for six days at high humidity.

### Preselection of fungal sRNA effector candidates

sRNAs either exclusively produced or increased in the *Mo*-*Bd* interaction were further investigated. In particular, differentially expressed 20-21 nt long sRNAs originating from non-coding regions of the *Mo* genome were considered potential sRNA effectors (ck-sRNAs) that target *Bd* genes as previously suggested for the *Botrytis cinerea* - *Arabidopsis thaliana*/*Solanum lycopersicum* pathosystems (Weiberg et al. 2013). Target prediction was carried out using psRNATarget with modified settings and a default score cut-off of 5.0. Some 3,691 fungal ck-sRNAs were predicted to target 45,066 *Bd* mRNAs in the necrotrophic phase of leaf infection (4 DPI), while fewer sRNA effector candidates (457 and 276, respectively) were predicted for the biotrophic phase (2 DPI) and the root setup corresponding to 24,077 and 16,083 mRNA targets (S2 Fig). Of note, the ratio between predicted targets and ck-sRNAs was different between biological samples, with the root sample having the highest (on average 58 predicted targets per ck-sRNA) compared to 53 and 12 for 2 DPI and 4 DPI leaf samples, respectively.

To substantiate a direct interaction of the predicted fungal sRNA effector candidates with *Bd* mRNAs during *Bd-Mo* interaction, we analyzed mRNA sequencing datasets from the same biological samples that were used for sRNA sequencing. This strategy allowed for the confirmation of target downregulation in presence of the predicted ck-sRNAs, which further selects forpotential sRNA effectors. As expected, many *Bd* and *Mo* genes were differentially expressed (up- or down-regulated) in the *Bd-Mo* interaction (Fig 6, S2 Tab), and a subset of these genes were differentially expressed in all three setups in roots and leaves, while others were tissue-specifically or fungal lifestyle-specifically (biotrophic, necrotrophic) induced (S3A-S3B Fig.). Overall, six *Bd* transcripts were found to be significantly (logFC < 0, padj < 0.05) downregulated in the biotrophic phase (2 DPI leaf samples), while 1,931 were downregulated in the necrotrophic phase (4 DPI leaf samples), and 38 in the *Mo*-infected root sample. Of these downregulated *Bd* transcripts, three were predicted to be plant targets of *Mo* ck-sRNAs in the 2 DPI sample, 1,895 in the 4 DPI sample, and eight in the root sample (Tab 1). In the next step, we assessed how many of these transcripts were targeted by sRNAs with 5’U, based on the consideration that Arabidopsis AtAGO1, which is involved in ckRNAi has a 5’ nucleotide preference (Weiberg et al. 2013, Mi et al. 2008). Following this strategy, we found two (leaf 2 DPI), 1,872 (leaf 4 DPI) and five (roots 4 DPI) potential *Bd* targets of fungal ck-RNAs in the different setups (Tab. 1). The predicted *Mo* sRNA / *Bd* mRNAs duplexes included genes for transcription factors such as *transcription factor MYB48-related* (BdiBd21-3.4G0132900.1) and *transcriptional regulator algH* (BdiBd21-3.1G0488800.1), exosome components (BdiBd21-3.4G0524000.1, BdiBd21-3.1G0012500.1, BdiBd21-3.1G0267100.1, BdiBd21-3.1G0357100.1, BdiBd21-3.4G0276900.1, BdiBd21-3.3G0350000.1), aquaporin transporters (BdiBd21-3.2G0400800.1, BdiBd21-3.3G0654800.1, BdiBd21-3.5G0207900.1, BdiBd21-3.5G0237900.1, BdiBd21-3.1G1005600.1), as well as RNA helicases, including the putative BdDCL3b (BdiBd21-3.2G0305700) (Tab. 2). A GO enrichment (GOE) analysis was carried out with AgriGO to detect over- and under- represented features in the dataset from the leaf 4 DPI setup, which covers the necrotrophic growth phase of *Mo*. In particular, generic GO terms associated with metabolic processes and photosynthesis were enriched (S4A-S4D Fig).

**Table 1.** Number of ck-sRNA effector candidates (20-21 nt) and their corresponding target mRNAs with significant (FC < 0, padj < 0.05) target downregulation.

| Type of ck-sRNA | Setup | No. of ck-sRNA with down-regulated targets | No. of 5' U ck-sRNA with down-regulated targets | No. of *Bd* / *Mo* targets | No. of *Bd* / *Mo* targets of 5'U ck-sRNAs |
| --- | --- | --- | --- | --- | --- |
| *Mo* ck-sRNAs | Leaf 2 DPI | 5 | 3 | 3 | 2 |
|  | Leaf 4 DPI | 3 436 | 2 546 | 1 895 | 1 872 |
|  | Root | 8 | 6 | 8 | 5 |
| *Bd* ck-sRNAs | Leaf 2 DPI | 954 | 186 | 907 | 423 |
|  | Leaf 4 DPI | 683 | 132 | 989 | 406 |
|  | Root | 697 | 124 | 237 | 125 |

Fold change (FC) and adjusted P value were determined by the analysis of mRNAseq data with DESeq2

**Table 2.**
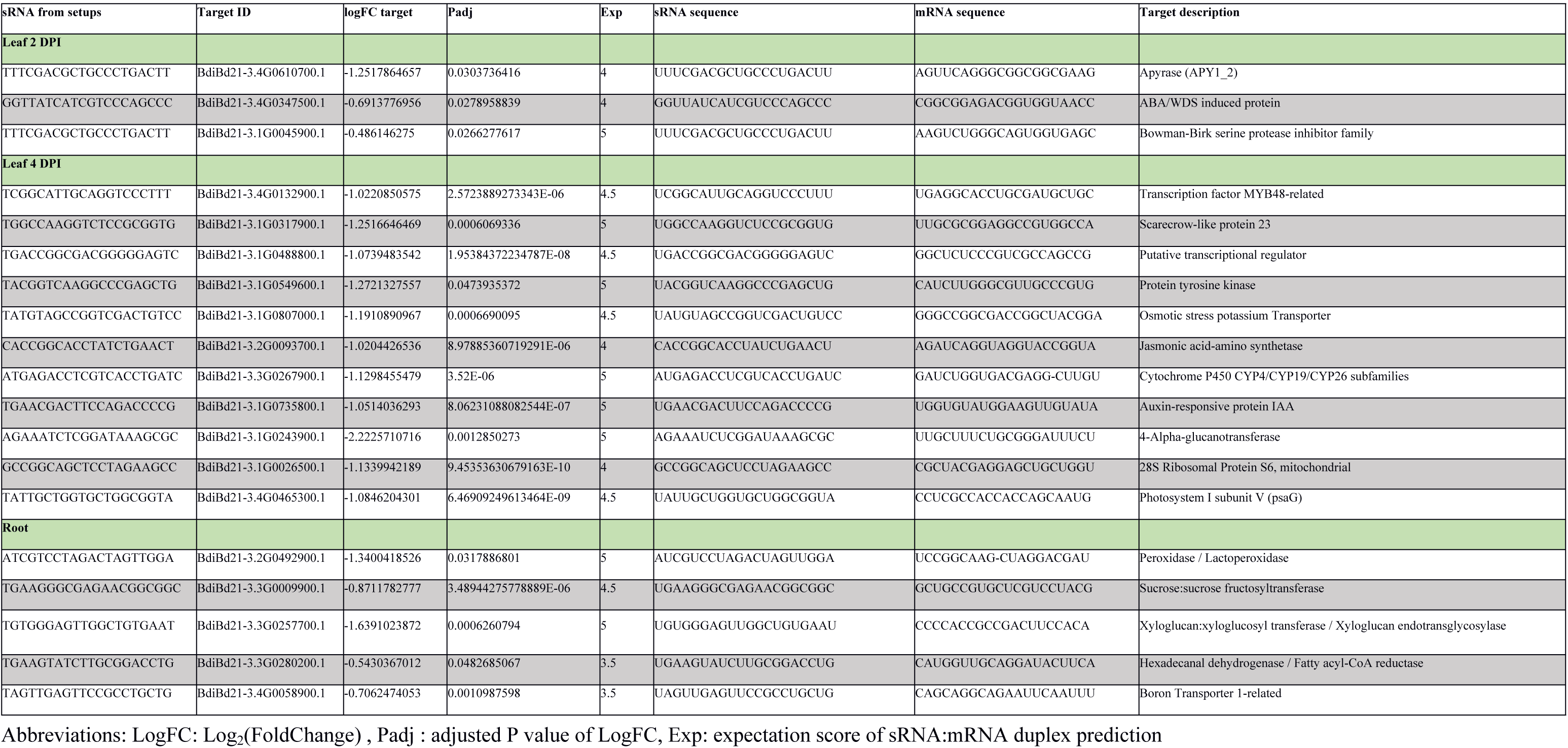
Selected *Mo* sRNA / *Bd* mRNA duplexes from infected *Bd* roots and leaves.

### Preselection of plant sRNA effector candidates

Given that plant-derived sRNAs have also been proven to move into fungal pathogens during plant colonization (Zhang et al. 2016; Cai et al. 2018b), we followed the same strategy used for the identification of candidate *Mo* ck-sRNAs to further analyze 20-21 nt sRNAs originating from the non-coding regions of the *Bd* genome showing a higher read count in the *Mo*-infected compared to non-infected samples. Target prediction for *Bd* sRNAs in the *Mo* transcriptome resulted in 1,070, 754 and 1,395 *Bd* ck-sRNA candidates for the 2 DPI and 4 DPI leaf and root setups, respectively (S5 Fig). The average number of *Mo* targets per *Bd* ck-sRNA candidate was relatively stable throughout the setups, with 7 to 12 targets predicted per *Bd* ck-sRNA. *Mo* mRNA levels were analyzed in both the infected samples and the axenic culture in order to substantiate the predicted target downregulation. *Mo* transcripts were significantly downregulated (logFC < 0, padj < 0.05) in the leaf 2 DPI (1,076), leaf 4 DPI (1,385) and in the root (287) setup (Fig 6, S2 Tab). Of these downregulated *Mo* mRNAs 907, 989 and 237, respectively, were predicted to be targeted by *Bd* ck-sRNAs (Tab.1). Focusing on those ck-sRNAs having 5’U, we further reduced the number of potential ck-RNAs and thus the number of potential *Mo* targets to 423 (leaf 2 DPI), 406 (leaf 4 DPI) and 125 (roots 4 DPI), respectively (Tab. 1). GOE analysis on the *Mo* mRNAs that were predicted as targets of *Bd* ck-sRNAs did not highlight significant differential representation in GO terms at 2 DPI and 4 DPI, while an enrichment in developmental (GO:0032502) and metabolic (GO:0008152) processes was detected in the root setup (S6A-S6E Fig). Confirmed downregulated *Mo* targets include cell wall related genes such as *chitin deacetylase 1* (*MGG_05023T0*), *chitinase 1* (*MGG_01247T0*), *cell wall protein MGG_09460T0* and virulence genes such as *CAP20* (*MGG_11916T0*) (Tab. 3). By comparing predicted fungal mRNA targets of *Bd* ck-RNAs in infected leaf and root tissue, we found a considerable overlap in significantly downregulated *Mo* targets between leaf and root samples (100 *Mo* mRNAs) and between the two leaf setups (354 *Mo* mRNAs), representing the biotrophic and necrotrophic phase of fungal colonization (Fig. 5, Fig 7).

**Figure 5.**
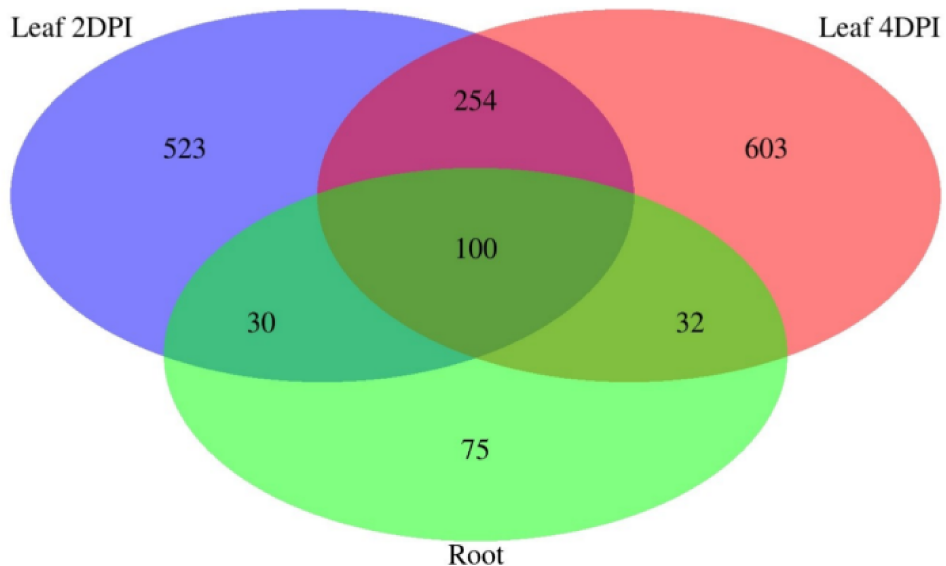
Venn diagram of downregulation *Mo* targets. Significantly downregulated (FC < 0 padj < 0.05) *Mo* mRNA targets with complementarity to *Bd* sRNAs shared between setups: leaf biotrophic phase (2 DPI; blue), leaf necrotrophic phase (4 DPI, red), and root (4 DPI, green). Transcript downregulation was assessed from mRNAseq data with DESeq2.

**Figure 6.**
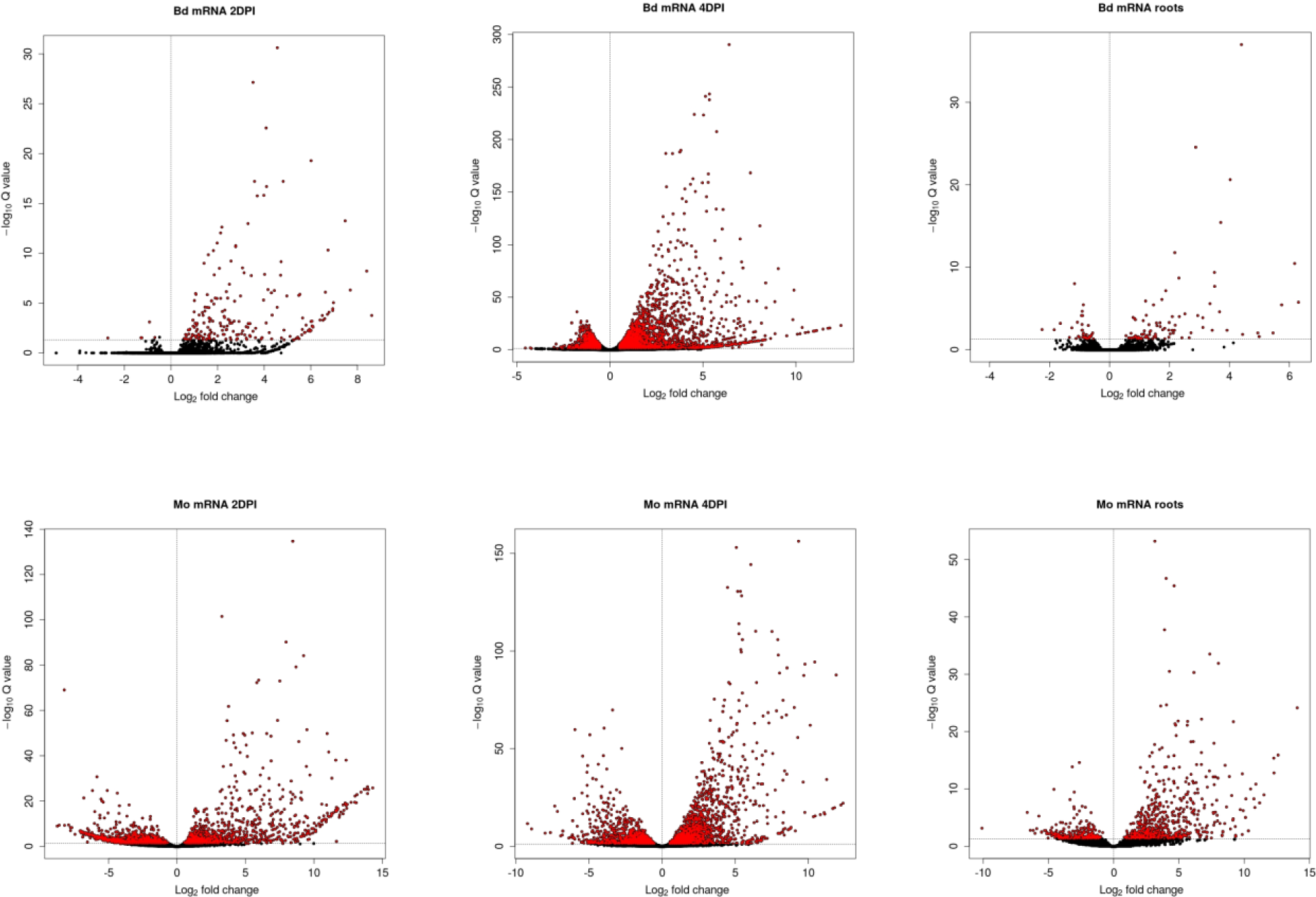
**(A,G) Volcano plots** of DESeq2 results for mRNAseq analysis of *Magnaporthe* and *Brachypodium* during infection.

**Figure 7.**
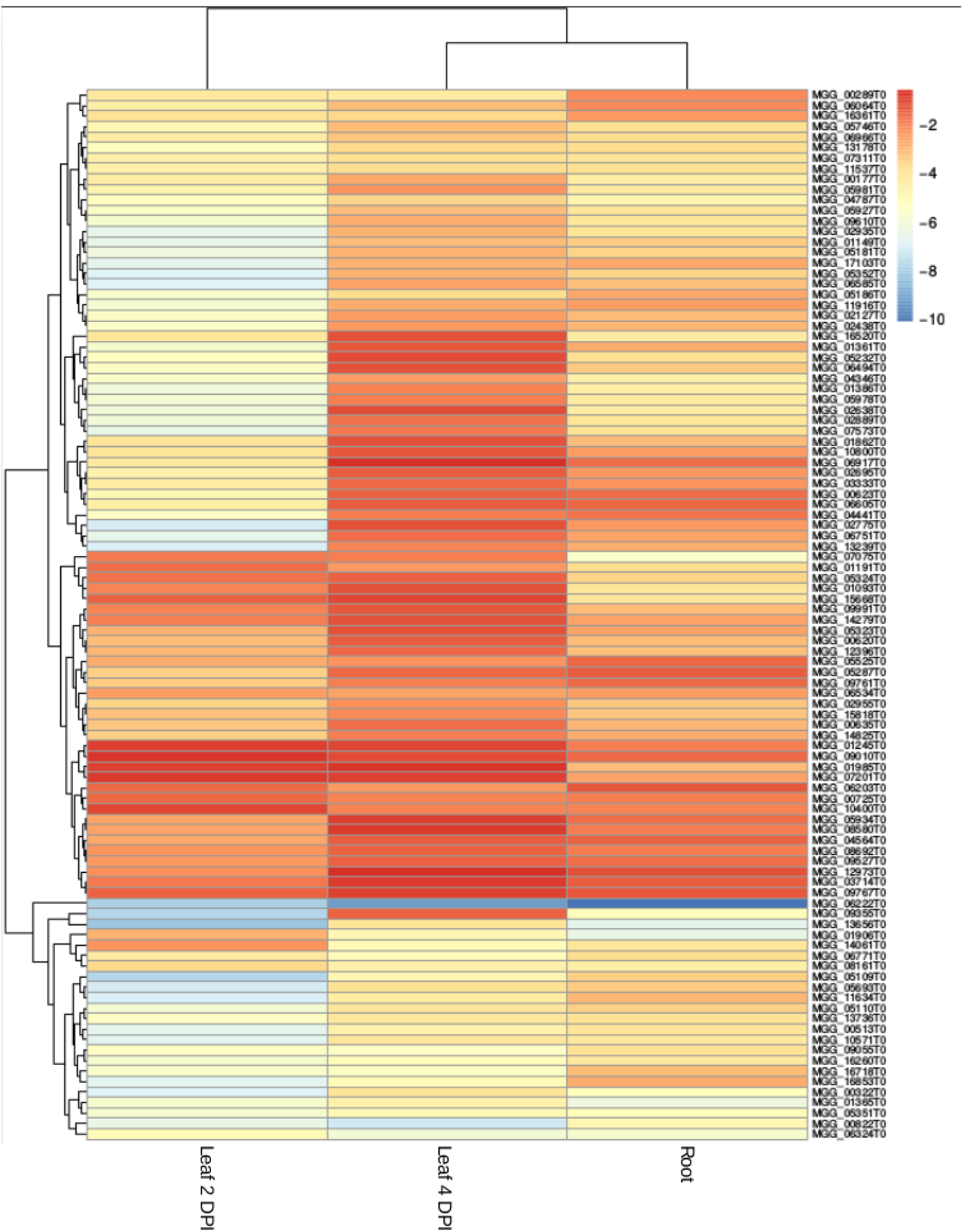
Heatmap for the *Mo* mRNA target expression levels. Heatmap of expression levels (logFC) of confirmed downregulated target *Mo* mRNAs in all 3 setups (leaf 2 DPI, leaf 4 DPI and root). Color gradient from red to blue indicative of log2FC of corresponding transcript (−0.5 (red) to −10 (blue)).

**Table 3.**
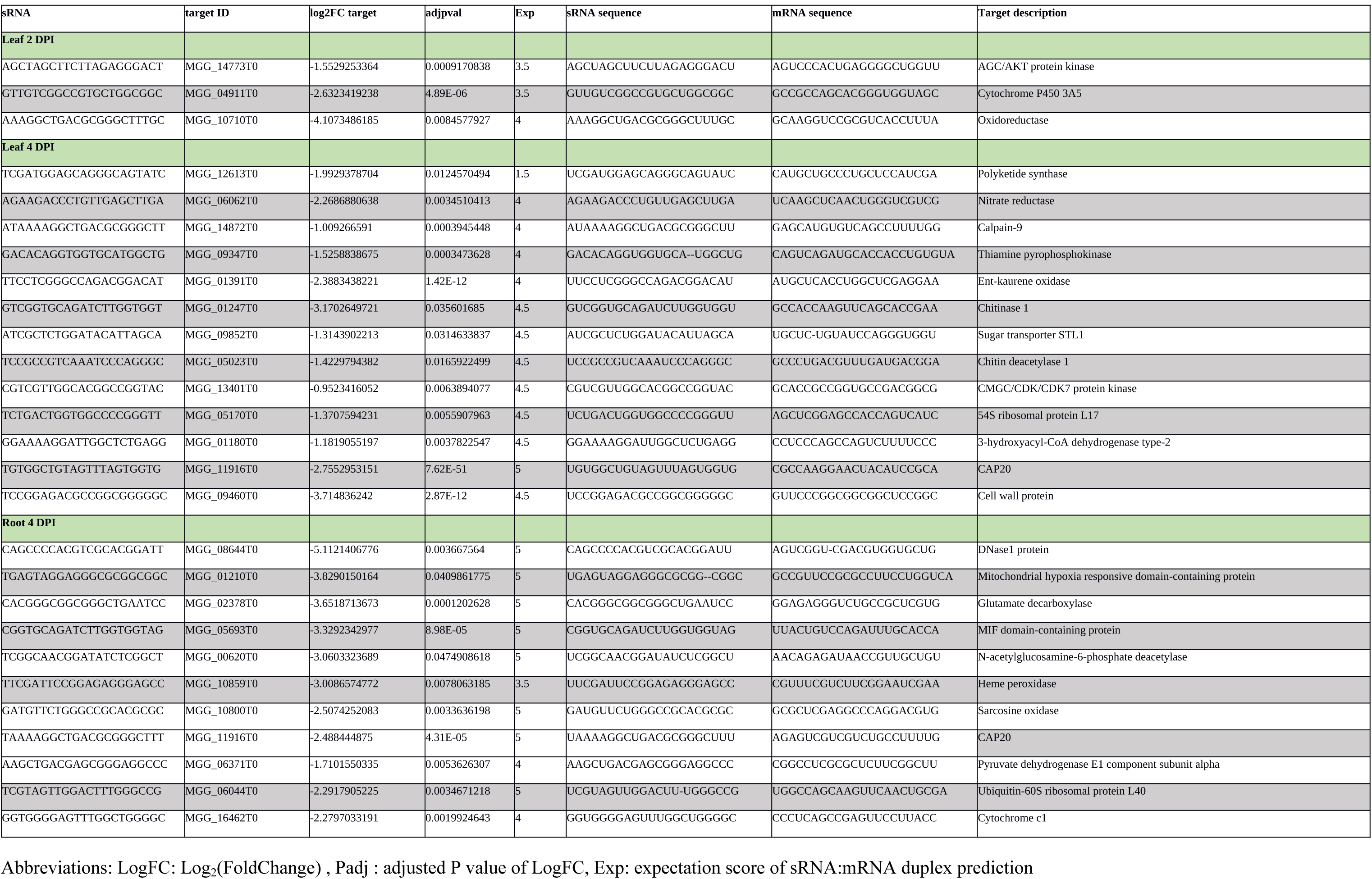
Selected *Bd* sRNA / *Mo* mRNA duplexes from infected *Bd* roots and leaves.

Next, we searched the PHI-base database for available information on the loss of virulence for the respective *Mo* target genes. A short list of down-regulated shared *Mo* mRNA targets and the PHI-base phenotypes are shown in (Tab. 4). Of note, we identified several *Mo* targets shared between the root and leaf setups that are known to be involved in *Mo* virulence and pathogenicity, including CON7 transcription factor (MGG_05287), the effector molecule AvrPiz-t (MGG_09055), N-acetylglucosamine-6-phosphate deacetylase (MGG_00620), chitin synthase D (MGG_06064), ATPase family AAA domain-containing protein 1 (MGG_07075), and mitochondrial DNA replication protein YHM2 (MGG_07201). Additionally, *Mo* mRNAs targets shared between the leaf infection timepoints included transcripts for autophagy-related protein MoATG17 (MGG_07667) and SNARE protein Sso1 (MGG_04090), whose respective mutants are also known to be compromised in pathogenicity (Kershaw and Talbot, 2009; Giraldo et al., 2013).

**Table 4.**
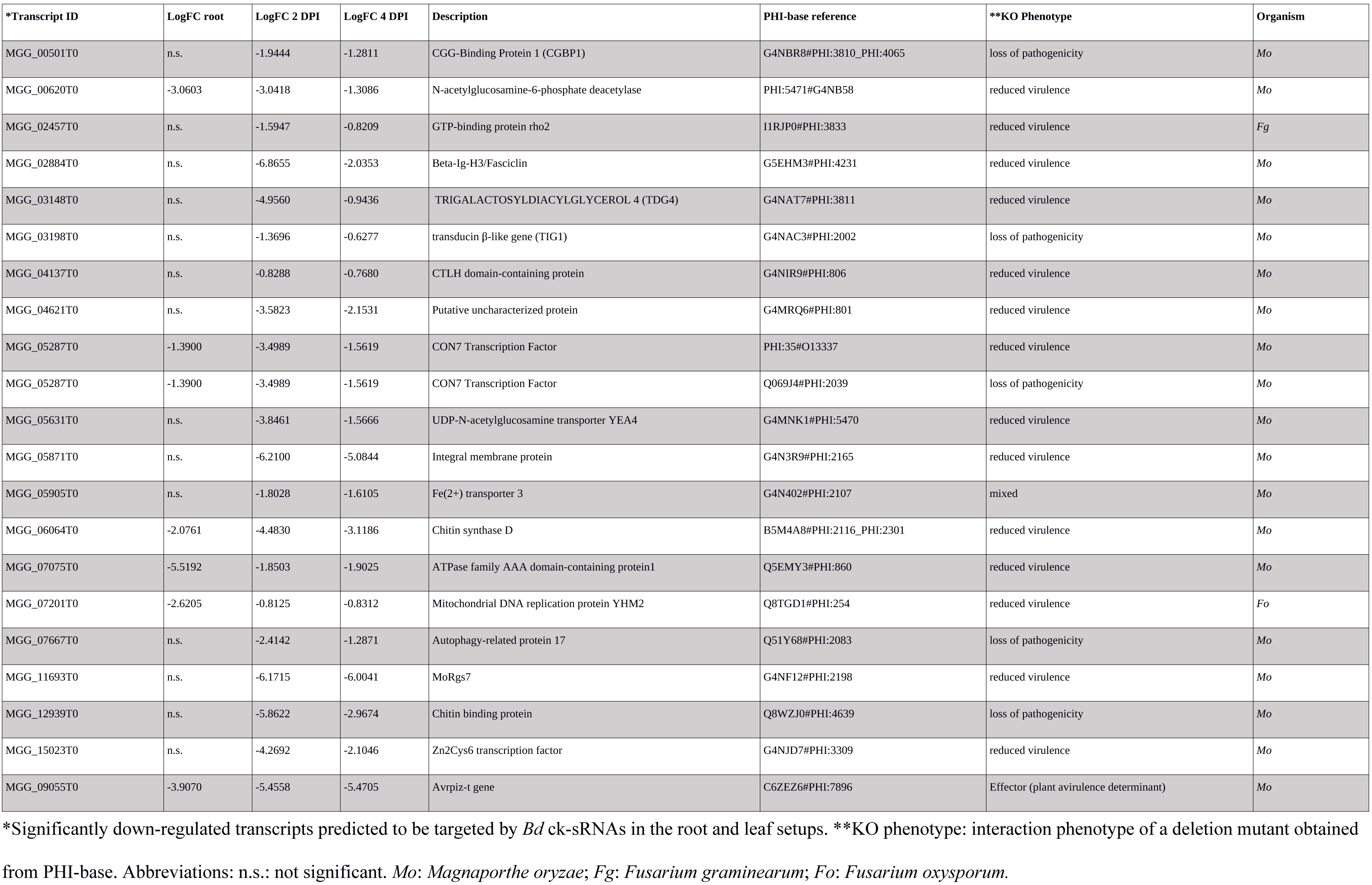
Selected shared Mo mRNAs targeted by Bd ck-sRNAs.

Overall, these results strongly suggest that *Bd* ck-sRNAs play a role in the defence response of the plant to rice blast infection and vice-versa, the fungus produces sRNA effectors to modulate Brachypodium metabolism and immunity.

## Discussion

In the present work we provide first experimental evidence for bidirectional RNA communication in the interaction of a monocotyledonous plant with its natural microbial pathogen. The *Brachypodium distachyon - Magnaporthe oryzae* pathosystem has been studied as a model for the blast disease of the staple crops rice and wheat, because *Bd* develops faster, has a smaller genome and requires less space for reproduction (Routledge et al. 2004; Parker et al. 2008; Vogel et al. 2006). Thus, our results support the possibility that major staple crops co-evolved mechanisms of RNA-based communication with their microbial pathogens. This notion is consistent with the important earlier observation that cereal plants are vastly amenable to biotechnological applications of dsRNA to control their pests and diseases (Koch et al. 2013; 2016; Cheng et al. 2015; Chen et al. 2016; Koch and Kogel 2014). The efficiency of HIGS, alike exogenous application of dsRNA (also called environmental RNAi or SIGS), requires both an operable RNAi machinery and a molecular basis for transfer of RNA between the interacting organisms (Koch et al. 2016; Wang et al. 2016). The detection of ckRNAi in *Bd* further substantiates the possibility that agronomic applications such as HIGS rely on evolutionarily evolved components and pathways for processing and transport of RNA. There have been reports that *Bd* employs RNAi in development and stress adaption: miRNAs have been proven to vary during exposure to abiotic stresses (Zhang et al. 2009) and between vegetative and reproductive tissues (Wei et al. 2009). Although the knowledge about the *Bd* RNAi machinery is less comprehensive, recent work predicted 16 AGOs and 6 DCLs, suggesting that the RNAi machinery is functional and follows the trend that cereals have extended families of key enzymes involved in RNAi (Mirzaei et al. 2014; Secic et al. 2019). *Magnaporthe oryzae* possesses a complete RNAi machinery and utilizes it throughout its development. *Mo* encodes two DCLs, three AGOs and three RdRPs (Kadotani et al. 2003; Murphy et al. 2008) and knock-out of RNAi pathway components severely affected the sRNA species produced by *Mo* and their accumulation levels *in vitro* (Raman et al. 2017). Moreover, sRNA-mediated alterations of TGS and PTGS have been detected *in vitro* both during starvation/different nutrient availability, and *in planta* during the different stages of rice leaf infection (Raman et al. 2013). Consistent with these observations, *Mo* mutants compromised in DCL and AGO function showed a reduced growth on *Bd21-3* leaves (Fig. 4). While a reduced virulence of *Δdcl1* and *Δdcl2* supports the hypothesis of ckRNAi as demonstrated earlier in the *B. cinerea* / Arabidopsis pathosystem (Weiberg et al. 2013), we cannot exclude though that the mutation in *MoDCL1* affects other processes that contribute to full virulence, which may also explain why mutations in AGO1 and AGO2 also reduced the fungal virulence. Moreover, we obtained all RNAi mutants from the D’Onofrio lab (Raman et al. 2013), and this group found no significant effects when the DCL mutants infected barley leaves. In our hands growth and development of said RNAi mutants was influenced strongly by growth conditions such as temperature and the culture medium. Considering this it can well be that the host genotype and/or growth media used for axenic cultures affects fungal development. Differences between sRNA libraries size distribution shown in Fig. 1 and the ones previously published by Raman et al (2013 and 2017) are due to the different protocols utilized both for sample preparation and for the data analysis itself. Those variations included: media utilized for fungal growth (OMA vs CM), inoculation protocol (drop inoculation onto Bd vs spray inoculation onto Os), sRNAs length selection (15-35 vs 20-30), sequencing machines and scripts for filtering sRNA reads applied during data analysis.

### Evidence for ckRNAi in the *Mo*-*Bd* interaction

To establish the origin of the sRNA reads detected in the different root and leaf setups of the *Mo-Bd* interaction, sRNAseq datasets from infected samples were aligned to both the *Bd 21-3* and the *Mo* 70-15 genome, and only reads aligning without mismatches to *Mo* and with at least two mismatches to *Bd* were assigned to the fungus and vice-versa, only reads aligning without mismatches to *Bd* and with at least two mismatches to *Mo* were assigned to the plant. As expected from the low amount of *Mo* in infected samples from leaves at the 2 DPI time point, most of the reads were assigned to *Bd*, whereas higher levels of *Mo* reads were detected in the 4 DPI leaf samples consistent with proliferating infection. All assigned reads were then filtered based on their read counts to select only reads either induced or upregulated in the datasets of infected tissues compared to uninfected tissues and axenic mycelia. We noted that most of the reads (>50%) found in infected samples are specific and are not detected in healthy tissues and axenic culture (Fig. 2), showing that sRNA production both in the plant host and the fungal pathogen is strongly responsive to infection. From this, it follows that sRNA datasets from healthy plants and axenic culture do not record the full diversity of sRNA communities. As an additional step we selected for sRNAs that were not aligning to the coding sequences of the organism of origin. The reasoning behind this filtering step is that we avoided accidental mRNA degradation to be kept as candidate sRNAs, and more important, we removed the sRNA sequences more likely to play an endogenous role (Zanini et al. 2018). Given that the size distribution of upregulated/induced sRNA reads did not show variation in peaks compared to the total sRNA reads (Fig. 1), we decided to select 21 nt sRNAs (canonical length for PTGS) and 20 nt sRNA (peak within the 20-24 nt sRNA population in *Mo*) for further analysis. Target prediction was carried out with psRNATarget, a web-based prediction software specifically designed for plant sRNA investigations. It allowed for the identification of complementary mRNA sequences in the interacting organism. Interestingly, we detected higher ratios of targets-to-sRNAs for *Mo* sRNAs in the leaf 2 DPI and the roots compared to the leaf 4 DPI sample, while *Bd* sRNAs showed lower and comparable averages.

In PTGS, sRNAs are loaded onto AGO proteins, which guide them towards a complementary mRNA sequence that will then be degraded or sequestered, resulting in reduced levels of the encoded protein. Knowing the expression levels of the predicted targets from the same biological samples used for the sRNA sequencing, we proceeded with further selection of candidate sRNAs based on the significant downregulation of their mRNA targets. Most of the predicted *Mo* ck-sRNAs in the 2 DPI leaf (biotrophic phase) and root samples did not pass this filtering step, as their predicted targets were either upregulated or had the same expression levels in the corresponding control datasets. There are a few possible explanations as to why the potential targets were not significantly downregulated in our mRNAseq datasets, including: i. the sRNA has not yet been transported throughout the tissue, so the downregulation is occurring only at the penetration site, where the fungus is physically interacting with the plant cells, and that is masked by the upregulation in distal parts of the tissue, ii. the target mRNA is not cleaved, but its translation is inhibited by the RISC complex acting as a physical barrier, in which case the measurable effect would not be at the mRNA level but only at the protein level, and iii. the target is indeed cleaved, but concurrently with the downregulating effect of the sRNA, there is a stronger endogenous upregulation of the gene, leading to either similar levels of mRNA as the control, or even higher. Importantly, in the 4 DPI leaf sample (necrotrophic phase), we observed that almost all significantly downregulated *Bd* mRNAs were predicted to have corresponding *Mo* sRNAs and the ratio of targets-to-sRNAs decreased from 12:1 predicted to 0.55:1 downregulated. Additionally, we checked the amount of confirmed *Mo* sRNAs that had a 5’U, known to be preferred by AtAGO1 for PTGS (Mi et al. 2008). We noted that 74% of the *Mo* sRNAs in the 4 DPI leaf sample had that base, and were predicted to target almost all (98.7%) the confirmed targets (Table 1).

### Fungal sRNA effectors

In order to substantiate the hypothesis that fungal ck-sRNAs function as effectors to aid the establishment and maintenance of infection, we investigated the role of putative downregulated *Bd* targets. Due to the low numbers of confirmed downregulated *Bd* targets in the 2 DPI leaf and root samples, we performed Gene Ontology Enrichment (GOE) only on the downregulated targets of the 4 DPI sample to assess whether specific functions or pathways were being targeted in *Bd* by *Mo* ck-sRNAs. Interestingly, GO terms associated with ribosomes and the photosystems were enriched in the target list compared to background, consistent with the hypothesis that *Mo* targets energy and metabolism of the plant to hinder its response to infection. Targeting conserved sequences such as ribosome- and photosynthesis-related ones would prove more efficient than specific defense / immunity genes that are more prone to mutate in the arms race between plants and pathogens. Specific plant targets included transcripts encoding for exosome components EXOSC1, EXOSC5, EXOSC6, EXOSC7, EXOSC8, EXOSC10 (Table 2). Extracellular vesicles have been recently discussed as the most likely mean of transport vehicle for ck-sRNAs and are in general known to cargo plant defense molecules to the infecting fungus (Rutter and Innes 2017; Baldrich et al. 2019; Cai et al. 2018b). Another subset of downregulated target transcripts included transcription factors such as members of the MYB family, PHOX2/ARIX, and the AP2/ERF family, known to regulate a multitude of cell processes, from plant development to hormone responses and biotic and abiotic stress responses (Ambawat et al. 2013; Cui et al. 2016). Interestingly, multiple *Brachypodium* aquaporin transporters (BdiBd21-3.2G0400800.1, BdiBd21-3.3G0654800.1, BdiBd21-3.5G0207900.1, BdiBd21-3.5G0237900.1, BdiBd21-3.1G1005600.1) were also effectively targeted by *Mo* sRNAs during the infection, consistent with the knowledge that aquaporins play a role in the interaction between plants and microbial pathogens, most likely by modulating both H_2_O availability and transport of reactive oxygen species (ROS; Afzal et al. 2016). Finally, a wide variety of genes involved in RNA metabolism was downregulated in *Bd*, from DNA-directed RNA polymerases subunits (RPB6, RPB12, RPC40) to RNA helicases, including the putative BdDCL3b (BdiBd21-3.2G0305700), involved in the preprocessing of sRNA precursor molecules involved in chromatin modification (Margis et al. 2006).

### Plant ck-sRNAs

We anticipated the plant to fight the spread of the infection by targeting vital/ virulence genes of *Mo*. To test this hypothesis, all confirmed downregulated *Mo* targets from leaves and roots were analyzed for gene ontology enrichment (GOE). While no relevant terms were found to be enriched or depleted in the 2 DPI and 4 DPI leaf samples, fungal metabolism and mycelia development related terms were enriched in the root target list, consistent with the aforementioned hypothesis. Comparison of *Mo* mRNA target lists between the different setups highlighted substantial target conservation between the leaf biotrophic and necrotrophic phases, with 354 shared *Mo* targets between the two, and 100 *Mo* targets conserved among all 3 setups. Subjecting the short list of *Mo* target genes to a PHI-base database survey for mutations in *Mo* with lethal or detrimental outcome, we found clear indication for a loss of virulence in respective KO mutants (Table 4). Additionally, among transcripts targeted at both leaf infection time points, we identified MoATG17 (MGG_07667T0) an autophagy-related protein, whose KO was previously shown to impair appressorium formation and function, resulting in a complete lack of disease symptoms on rice leaves (Kershaw and Talbot, 2009). Moreover we found t-SNARE Sso1 (MGG_04090T0) previously proven to be involved in the accumulation of fungal effector molecules at the biotrophic interfacial complex (BIC) during rice leaf infection (Giraldo et al. 2013).

Common targets between all three setups (leaves and root infection alike) included various fungal cell wall related genes, namely acidic endochitinase SE2 (MGG_03599), chitinase (MGG_04534), GPI-anchored cell wall beta-1,3-endoglucanase EglC (MGG_10400), and chitin synthase D (MGG_06064). Interestingly, genes known to be involved in the maintenance of the disease were also targeted. For instance, CON7 transcription factor (MGG_05287), known to regulate the expression of a wide range of infection-related genes (Shi et al. 1995; Odenbach et al. 2007), is targeted and significantly downregulated across all infection datasets. Additionally, we detected sRNAs targeting the mRNA encoding for the avirulence effector molecule AvrPiz-t (MGG_09055). AvrPiz-t suppresses rice PTI signaling pathway by targeting the E3 ubiquitin ligase APIP6 and suppressing its ligase activity, resulting in reduced flg22-induced ROS generation and overall enhanced susceptibility *in vivo* (Park et al., 2012).

## Conclusions

Taken together our results provide the first experimental evidence of bidirectional cross-kingdom RNAi within a monocot pathosystem, and strongly support the model that sRNAs play a crucial role in ckRNAi during plant host - pathogen interactions, including systems of staple field crops. Furthermore, ck-RNAs induced during infections show only partial overlap both among the different tissues (leaves, roots) and the different infection phases (leaf: biotrophic, necrotrophic), showing that ckRNAi in a given host - pathogen interaction exhibits tissue- and lifestyle-specificity.

## Material and Methods

### Sample preparation from *Mo*-*Bd* interactions

*Magnaporthe oryzae* (*Mo* 70-15; Raman et al. 2013) was grown on oatmeal agar (OMA) for two weeks at 26°C with 16 h light/8 h dark cycles both for sampling of mycelium and conidia production. Samples from axenic cultures were collected by scraping a mixture of mycelia and spores from three plates, followed by immediate freezing in liquid nitrogen. For root inoculation, sterilized seeds of *Brachypodium distachyon* genotype *Bd21-3* (Vogel & Hill, 2008) were vernalized in the dark at 4°C for two days on half strength MS (Murashige and Skoog 1962) medium and then moved to a 16 h light/8 h dark cycle at 22°C/18°C. Roots of one-week-old seedlings were dip-inoculated in 1 ml of conidia solution (250,000 conidia/ml in 0.002% Tween water) for 3 h, transplanted in a (2:1) mixture of vermiculite (Deutsche Vermiculite GmbH) and Oil-Dri (Damolin, Mettmann, Germany) and grown for additional 4 days before harvesting. Control roots were mock-inoculated with 1 ml of Tween water solution. For leaf inoculation, third leaves of three-week-old *Bd21-3* were detached and drop-inoculated with 10 μl of conidia solution (50,000 conidia/ml in 0.002% Tween water) on 1% agar plates. Control leaves were mock-inoculated with Tween water. Leaves were collected for sequencing at 2 DPI (days post inoculation) and 4 DPI. *Mo* 70-15 mutants *M. oryzae Δmoago1*, *Δmoago2*, *Δmoago3, Δmodcl1, Δmodcl2, Δmodcl1/2 and Δmodcl2/1*, obtained from N. Donofrio, Newark, U.S.A, were grown and inoculated onto *Bd21-3* leaves as described above, with the exception of *Δmoago3* that failed to sporulate and was not further tested. *Mo* lesions were assessed at 6 DPI.

### RNA extraction, library preparation and sequencing

Three roots or two leaves, respectively, were pooled per sample for RNA extraction and for each condition three pooled biological samples were prepared. Frozen tissue stored at −80°C was ground in liquid nitrogen using mortar and pestle. Total RNA was isolated with ZymoBIOMICS TM RNA Mini Kit (Zymo Research, USA) according to the manufacturer’s instructions. Quantity and integrity of the RNA were assessed with DropSense16/Xpose (BIOKÉ, Netherlands) and Bioanalyzer 2100 (Agilent, Germany), respectively. Purification of small and large RNAs into separate fractions was carried out using RNA Clean & Concentrator TM −5 (Zymo Research) and concentration and quality of the fractions were checked again. Fifty ng of small RNA (17 to 200 nt) were used for cDNA library preparation with TruSeq® Small RNA Library Prep (Illumina, USA) and 1.5 μg of large RNA were used for cDNA library preparation with TruSeq^®^ Stranded mRNA (Illumina). Constructed cDNA libraries of sRNAs were further size selected with BluePippin (Sage Science, USA) for fragments between 140 and 160 nt in length (15-35 nt without adapters). Quality of polyA mRNA libraries was assessed using the Fragment AnalyzerTM Automated CE System (Advanced Analytical Technologies, Austria).

The Illumina HiSeq1500 sequencing platform was used to sequence the Illumina TruSeq^®^ Small RNA libraries single end with 35 nt read length and the Illumina TruSeq^®^ Stranded mRNA libraries (paired-end [PE] sequencing, 70 nt) of all samples.

### sRNA analysis

The single end sequenced cDNA reads of Illumina TruSeq^®^ Small RNA libraries were analyzed starting with quality check with FastQC (Andrews 2010) and trimming of adapter artifacts with cutadapt (Martin 2011). The alignment of the reads to reference genomes and transcriptomes of *Bd* and *Mo* was done using the short read aligner Bowtie (Langmead et al. 2009). Reads with a 100% alignment to the genome of the organism of origin were selected, alongside the reads with at least two mismatches in the alignment to the target organism genome.

### Identification of sRNA effectors

Bioinformatics analysis of sRNAs effectors was done as described (Zanini et al. 2018). Only sRNA reads of 20-21 nt length originating from non-coding regions and with a higher count in the organism of origin control datasets compared to the infected ones were analyzed further for sRNA effector identification by the target prediction software psRNATarget used with customized settings (Dai & Zhao 2011).

### mRNA analysis and sRNA target confirmation

Paired end sequenced cDNA reads of Illumina TruSeq^®^ Stranded mRNA libraries were analyzed through the quality check in FastQC and alignment in the junction mapper HISAT2 (Kim et al. 2015). Htseq-count (Anders et al. 2014) and DESeq2 (Love et al. 2014) were then used for differential expression gene calling (DEG) between the infected and control sample genes. Expression levels obtained for each gene were used as confirmation of downregulation of predicted targets from the psRNATarget software. Gene Ontology Enrichment analysis on the confirmed targets was carried out with Agrigo (Du et al. 2010). PHI-base, a collection of experimentally verified pathogenicity/virulence genes from fungal and microbial pathogens (Baldwin et al. 2006), was used to gather information regarding phenotype and virulence of fungal mutants carrying a mutation in the identified *Mo* gene targets.

## Author Contributions Statement

KHK, SZ, ES, TB andJK and designed the experiments; SZ, ES, TB and JK conducted the bioinformatics analysis; SZ and KHK wrote the text.

## Acknowledgment

We thank Elke Stein, Dagmar Biedenkopf, and Christina Birkenstock for technical assistance. We thank Dr. John Vogel and the DOE-JGI for permission to use the Bd21-3 genome under early access conditions. We are grateful to Nicole M. Donofrio, Department of Plant & Soil Sciences, University of Delaware, Newark, for sharing the *Magnaporthe oryzae* mutants. *Brachypodium distachyon Bd21-3* is a gift of R. Sibout, INRA Verseille.

## Supplemental Data

**S1 Fig.** Size distribution of total filtered reads in the interaction of *Brachypodium distachyon* and *Magnaporthe oryzae*.

**S2 Fig.** Schematic overview of *Mo* sRNAs effectors (20-21 nt) and corresponding *Bd* target mRNAs after target prediction with psRNATarget with customized settings.

**S3 Fig. A-B** Selected differentially expressed *Bd* and *Mo* transcripts

**S4 Fig. A-D** Results of gene ontology enrichment (GOE) analysis for significantly downregulated *Bd* mRNA targets in the 4 DPI leaf setup. GOE analysis done with AgriGO.

**S5 Fig.** Overview of *Bd* sRNAs (20-21nt) effectors and corresponding *Mo* mRNAs after target prediction with psRNATarget with customized settings.

**S6 Fig. A-E** Overview of gene ontology enrichment (GOE) analysis for significantly downregulated *Mo* mRNA targeted in the root setup. GOE analysis done with AgriGO.

**S1 Tab.** Overview of total sRNA and mRNA reads in the *Brachypodium distachyon* – *Magnaporthe oryzae* interaction.

**S2 Tab.** Total numbers of significantly (padj < 0.05) up- or down-regulated genes in the *Brachypodium distachyon* – *Magnaporthe oryzae* interaction (DESeq2 results)

